# Rapid Automated Large Structural Variation Detection in a Diploid Genome by NanoChannel Based Next-Generation Mapping

**DOI:** 10.1101/102764

**Authors:** Alex R Hastie, Ernest T Lam, Andy Wing Chun Pang, Xinyue Zhang, Warren Andrews, Joyce Lee, Tiffany Y Liang, Jian Wang, Xiang Zhou, Zhanyang Zhu, Thomas Anantharaman, Željko Džakula, Sven Bocklandt, Urvashi Surti, Michael Saghbini, Michael D. Austin, Mark Borodkin, R. Erik Holmlin, Han Cao

## Abstract

The human genome is diploid with one haploid genome inherited from the maternal and one from the paternal lineage. Within each haploid genome, large structural variants such as deletions, duplications, inversions, and translocations are extensively present and many are known to affect biological functions and cause disease. The ultimate goal is to resolve these large complex structural variants (SVs) and place them in the correct haploid genome with correct location, orientation, and copy number. Current methods such as karyotyping, chromosomal microarray (CMA), PCR-based tests, and next-generation sequencing fail to reach this goal either due to limited resolution, low throughput, or short read length.

Bionano Genomics’ next-generation mapping (NGM) offers a high-throughput, genome-wide method able to detect SVs of one kilobase pairs (kbp) and up. By imaging extremely long genomic molecules of up to megabases in size, the structure and copy number of complex regions of the genome including interspersed and long tandem repeats can be elucidated in their native form without inference.

Here we tested Bionano’s SV high sensitivity discovery algorithm, Bionano Solve 3.0, on *in silico* generated diploid genomes with artificially incorporated SVs based on the reference genome, hg19, achieving over 90% overall detection sensitivity for heterozygous SVs larger than 1 kbp. Next, in order to benchmark large SV detection sensitivity and accuracy on real biological data, we used Bionano NGM to map two naturally occurring hydatidiform mole cell lines, CHM1 and CHM13, each containing a different duplicated haploid genome. By *de novo* assembling each of two mole’s genomes separately, followed by assembling a mixture of CHM1 and CHM13 data, we were able to measure heterozygous SV sensitivity by comparing SVs called in the mixture assembly against those called in the individual assemblies. We called 1999 unique SVs (> 1.5 kbp) in the pseudo-diploid assembly and established 87.4% sensitivity for detection of heterozygous SVs and 99.2% sensitivity for homozygous SVs. In comparison, a recent SV study on the same CHM1/CHM13 samples using long read NGS alone showed 54% sensitivity for detection of heterozygous SVs and 77.9% for homozygous SVs larger than 1.5 kbp. We also compared an SV call set of the diploid cell line NA12878 with the results of an earlier mapping study (Mak AC, 2016) and found concordance with 89% of the detected SVs found in the previous study and, in addition, 2599 novel SVs were detected. Finally, two pathogenic SVs were found in cell lines from individuals with developmental disorders. *De novo* comprehensive SV discovery by Bionano NGM is shown to be a fast, inexpensive, and robust method, now with an automated informatics workflow.

## Introduction

The importance of understanding the role of large SVs in genetic disorders cannot be overstated. For many known syndromes, clinically relevant large SVs are well characterized: deletions have been found to cause Prader-Willi syndrome, DiGeorge syndrome, and Williams-Beuren syndrome; duplications can cause Charcot-Marie-Tooth disease, and inversions can cause haemophilia A (Emanuel BS, 2001). More recently, large SVs have been found to play a role in neurological disease, such as early-onset neuropsychiatric disorders (Brand H, 2014), Tourette syndrome (Fernandez TV, 2012), and Parkinson’s disease (Butcher NJ, 2013); and in coronary heart disease (Crawford DC, 2008) and congenital heart disease (Bittel DC, 2014). SVs have also been shown to influence obesity (Bochukova EG, 2010) and pharmacogenetics (Rasmussen HB, 2012).

Cancer cells typically show extreme rearrangements of the genome. SVs are found in most cancer types. Examples are FGFR3-IGH fusion genes in multiple myeloma caused by a translocation and deletion, or the Philadelphia chromosome found in chronic myeloid leukemia (Krem MM, 2015).

Existing technologies including chromosomal microarrays (CMA) and whole genome sequencing diagnose fewer than 50% of patients with suspected genetic conditions (Miller DT, 2010) (Lee H, 2014). This leaves a majority of patients without ever receiving a molecular diagnosis. Chromosomal microarray lacks the ability to detect structural variants in which no loss or gain of sequence took place. Balanced translocations and inversions are essentially invisible to these technologies.

Clinical exome sequencing solves about 30% of rare diseases (Lee H, 2014). NGS reliably identifies single nucleotide variants and small insertions and deletions. It relies on short-read sequences that are mapped to a reference human genome and has limited power to identify most larger insertions, deletions, and copy-number variations. Various NGS based SV calling algorithms routinely disagree with each other and have limited power to detect SVs such as inversions and translocations (Alkan C, 2011). Huddleston *et al.* used long read technology by Pacific Biosciences to call structural variants in cell lines that were extensively analyzed using short-read sequencing as part of the 1000 Genomes Project. They report that > 89% of the variants identified using long reads have been missed as part of the analysis of the 1000 Genomes Project, even after adjusting for more common variants (Huddleston, 2016).

The true extent of structural variation of the genome has been enigmatic as many types of structural variation are refractory to current technologies. Estimates suggest that structurally variable regions cover 13% of the human genome, and individuals show structural variation covering as much as 30 Mbp between each other (Sudmant PH, 2016). This makes large structural variation the biggest source of individual genomic variation (Pang, 2010). In addition, the human genome is diploid with one of each haploid genome inherited from the maternal and the paternal lineage. Phenotypical or pathological outcomes are results of interplay between variants of the two haploid genomes. In order to understand genetic diseases and develop treatments and diagnostic tools for them, it is important to be able to identify these SVs and determine which variant belongs to which haploid genome. To completely resolve these large complex structural variants such as balanced chromosomal lesions and interspersed repeats and place them in their appropriate haploid with correct location, orientation and copy number has been the ultimate goal in genomic research and precision diagnostics. Current methods such as karyotyping, CMA, PCR-based tests, and next-generation sequencing fail to reach this goal, either due to their limited resolution, low throughput, or short read length.

Bionano Genomics’ Next-Generation Mapping (NGM) can elucidate all SV types, larger than 1 kbp. Extremely long (hundreds of kilobases to multiple megabases) genomic DNA is extracted from the sample source and fluorescently labeled at nicking endonuclease recognition sites consisting of specific 6-7 bp sequences, after which it is linearized and uniformly stretched in high density NanoChannel arrays, and imaged on the Irys System (Lam et al.,). Since the molecules are linear and uniformly stretched, the distances between labels can be accurately measured, resulting in single molecule maps with a unique labeling pattern (“barcode”) which has a density of approximately 10 times per 100 kbp on average (for human genomes). Using an overlap-layout-consensus assembly algorithm, consensus genome maps are constructed, refined, extended, and merged to create a *de novo* diploid genome map assembly useful for heterozygous SV detection. Multiple genome map assemblies can be created, each using a different endonuclease, and may be scaffolded together along with the reference sequence to generate broader coverage and higher label density.

Bionano maps are built completely *de novo*, based on specific nick-labeling patterns on native long intact molecules, without any reference guidance. This differentiates NGM from NGS, or inferred synthetic long reads, where short-read sequences are typically aligned to a reference and reconstructed. When short reads are forced to align to a potentially incorrect, incomplete or divergent reference, or when mismatched reads are excluded from the alignment, SVs in the region may be missed or mischaracterized. More importantly, short-read based technologies are less likely to readily detect novel insertions, mobile element insertions, variable number tandem repeat variants or balanced SVs such as inversions and translocations such as those that cause certain diseases. *De novo* constructed individual genomes using Bionano maps allow for comprehensive structural variation analysis.

## Results

We report the use of a fully automated SV calling pipeline for Bionano NGM to identify large (> 1 kbp) homozygous and heterozygous SVs with unprecedented sensitivity and specificity.

To benchmark large SV detection sensitivity and accuracy in a diploid genome, we first generated an *in silico* map of the human reference genome, hg19, and incorporated random SVs to create an *in silico* diploid genome. We used this to test Bionano’s sensitivity and specificity to detect insertions, deletions, and translocations against the *in silico* “ground truth.”

Next, we used Bionano NGM to map two naturally occurring hydatidiform mole cell lines, CHM1 and CHM13, each containing a different duplicated haploid genome. By *de novo* assembling each of two mole genomes separately, followed by assembling a mixture of CHM1 and CHM13 molecules, we were able to compare the simulated diploid, heterozygous SV calls against the homozygous SV calls in each single haploid mole assembly.

We further tested this new pipeline in a well characterized diploid cell line NA12878, which was previously mapped with Bionano Irys technology and heavily manually curated (Mak AC, 2016). The overall SV detection sensitivity improved more than fourfold, primarily by improving sensitivity for insertions and deletions down to 1 kbp from the 5 kbp threshold set in the previous study. Finally, the Bionano SV workflow was applied to cell lines containing known translocations, GM16736 and GM21891.

### Simulation

Translocations were simulated similarly to insertions and deletions (Table 1). There were 4 diploid genomes containing 1,844 (±17) cut-and-paste translocations, and 3 diploid genomes containing 48, 6, and 6 reciprocal translocations. Details of the cut-and-paste and reciprocal translocation simulation are described in the Methods section.

**Table 1.**
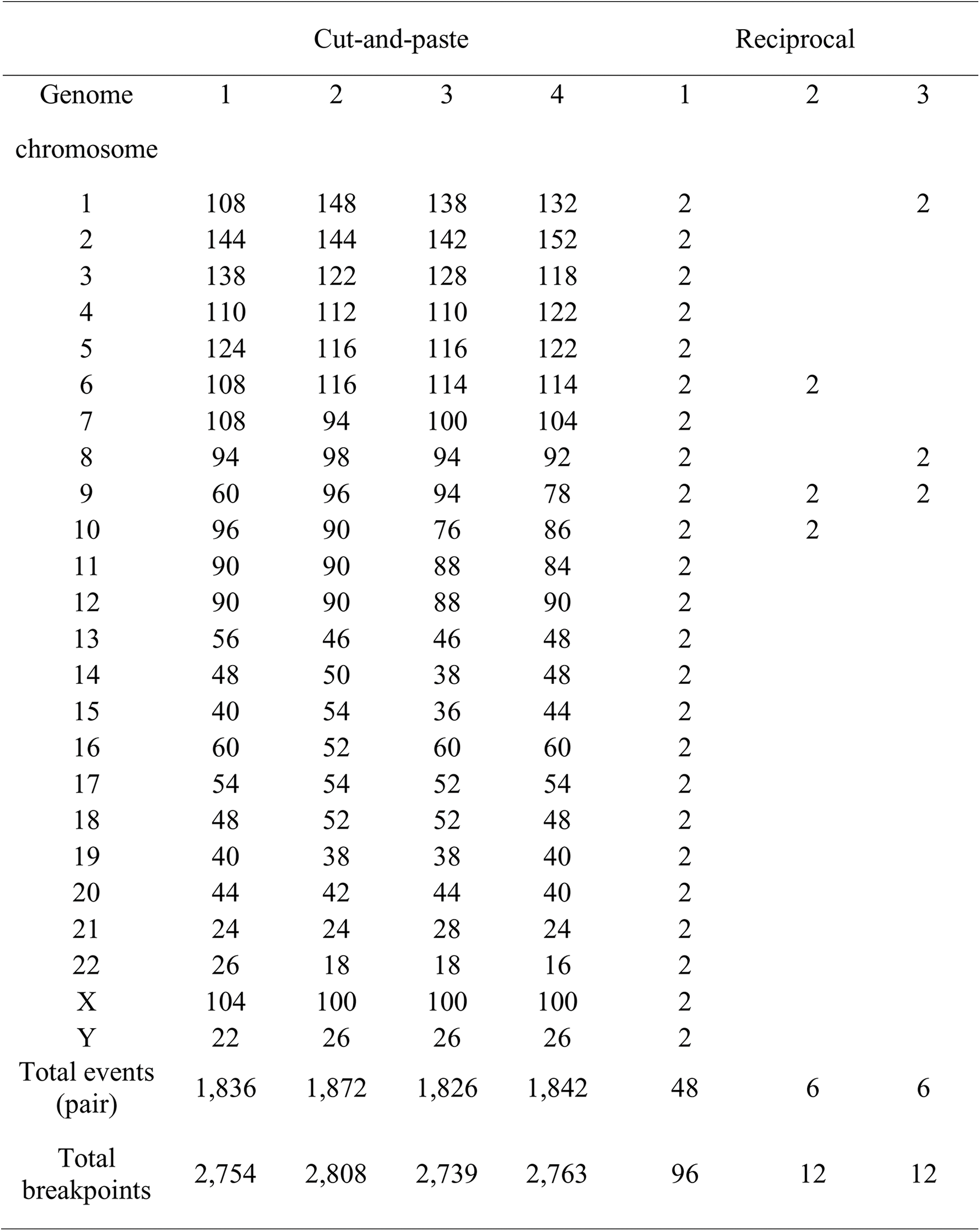
Number of simulated translocation events along hg19. The events were calculated as pair of breakpoints. Translocations on cut-and-paste genomes and reciprocal genome 1 were randomly simulated. Translocations on reciprocal genomes 2 and 3 were from reference studies.

### *In silico* insertion, deletion, and translocation sensitivity estimates

Figure 1 shows sensitivity and positive predicted value (PPV) for heterozygous insertions and deletions within a large size range. SVs calls were generated independently for each of the two nicking enzymes, and sensitivity and PPV are plotted for each as well as a combined SV set (SVMerge), which is a result of combining unique SVs from each enzyme and collapsing SVs called by each enzyme (merged SVs also have higher confidence as they are called independently twice). Overall sensitivity for heterozygous SVs larger than 1 kbp was 90.2% for SVMerged SVs. Insertion and deletion size measurements have only a 49 bp median error, while reported breakpoints were a median distance of 3.3 kbp from the actual breakpoint coordinates. Breakpoint resolution is limited by the density of nick motif (every 5 kb on average when both Nt.BspQI and Nb.BssSI enzymes are used). Sensitivity seemingly decreases for large (>200 kbp) insertions due to the simulation of randomly inserted fragile sites. Some of these insertions appear as “end” SVs if they are sufficiently large.

**Figure 1:**
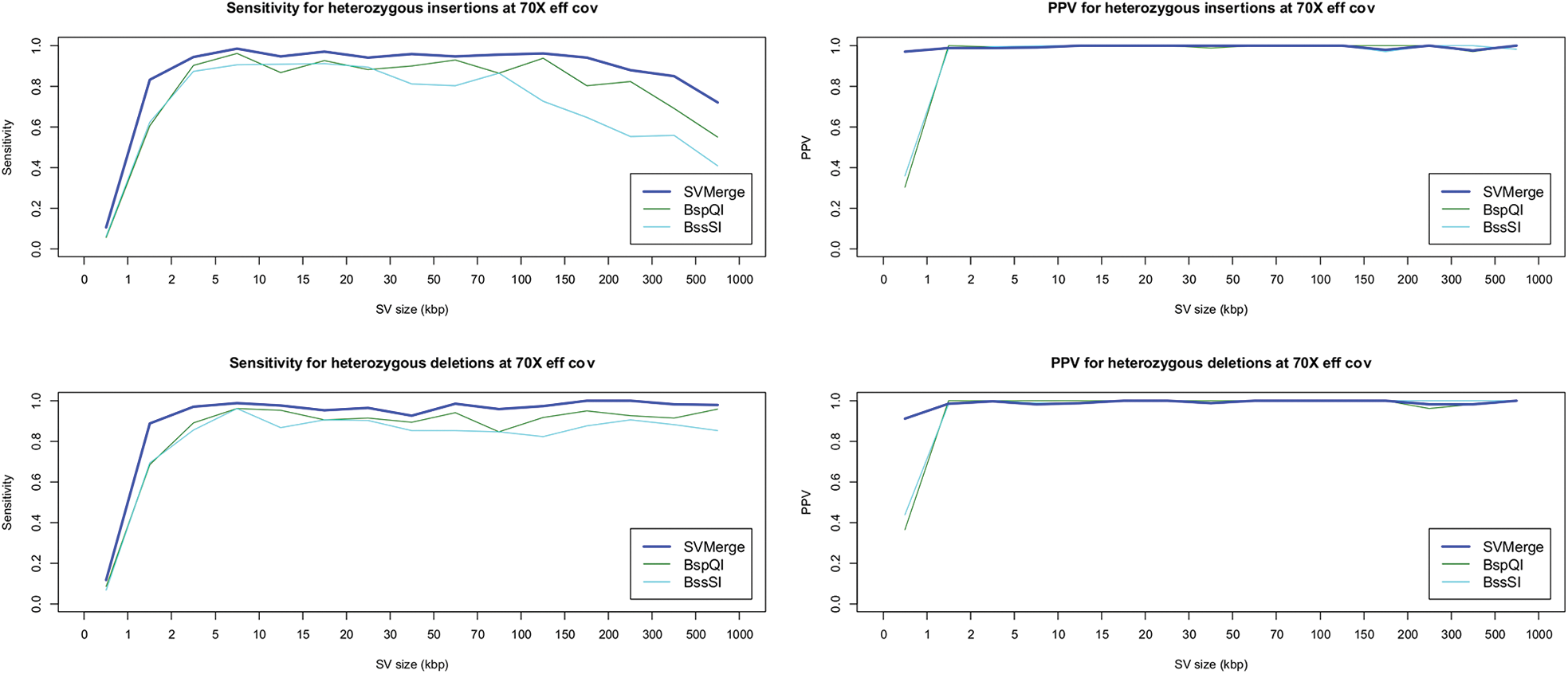
Heterozygous SV calling performance from a simulated dataset. Molecules were simulated from unedited and edited versions of hg19 (with insertions and deletions of different sizes) and used for assembly and SV calling. SV detection sensitivity and PPV across different size ranges are shown based on assemblies from molecules labeled with nicking endonuclease Nt.BspQI, Nb.BssSI, and combined data from both motifs (SVMerge).

Sensitivity for heterozygous translocations was 98% for breakpoint detection after SVMerge; single enzyme sensitivities were 83% (Nb.BssSI) and 90.7% (Nt.BspQI). Because NGM has relatively high precision relative to cytogenetic assays, many breakpoints are defined to within a few kbp of the precise breakpoint, and this distance is often close enough for PCR validation of translocations discovered by NGM. Genome mapping can define the true positions of breakpoints within a median distance of 2.9 kbp.

### Homozygous and heterozygous SVs in a pseudo-diploid mixture genome based on CHM1 and CHM13 double haploid cell lines

Since there is no structurally accurate haploid resolved genome sequence available from a natural diploid human sample that can be used as the ground truth to evaluate a technology and algorithm for sensitivity in detecting SVs, a diploid human genome was created by combining data from two haploid hydatidiform mole derived cell lines, as was done by Huddleston *et al*. These moles usually occur when an oocyte without nuclear DNA gets fertilized by a sperm. The haploid genome in the sperm is duplicated, and the cell line resulting from this tissue (such as CHM1 and CHM13) is therefore expected to be entirely homozygous.

Genomic DNA from CHM1 and CHM13 was isolated, labeled at Nt.BspQI nick sites and at Nb.BssSI nick sites in separate reactions. Molecules imaged in each cell line were *de novo* assembled using the Bionano Solve 3.0 pipeline. SVs detected in the homozygous cell lines were considered the (conditional) ground truth. An equal mixture of randomly selected single molecule data from the two cell lines was assembled to simulate a diploid genome, and SV calls made from this mixture were used to estimate the pipeline’s sensitivity to detect homozygous and heterozygous SVs. Assembly statistics are shown in Table 2.

**Table 2:** Metrics for assemblies from data from CHM1, CHM13, and an equal mixture of both. Samples were labeled at Nt.BspQI and at Nb.BssSI nick sites in separate reactions.

| Sample | CHM1 |  | CHM13 |  | Mixture |  |
| --- | --- | --- | --- | --- | --- | --- |
|  | Nt.BspQI | Nb.BssSI | Nt.BspQI | Nb.BssSI | Nt.BspQI | Nb.BssSI |
| Data collected (Gbp)<br>for molecules >150 kbp | 251.32 | 219.08 | 315.78 | 341.30 | 374.90 | 344.54 |
| Molecules N50 (kbp) | 277.99 | 239.63 | 308.60 | 289.79 | 292.60 | 246.46 |
| Assembly size (Gbp) | 2.90 | 2.89 | 2.90 | 2.87 | 5.63 | 5.76 |
| Genome map N50<br>(Mbp) | 7.07 | 2.61 | 4.48 | 2.18 | 7.10 | 3.00 |

Table 3 shows the number of insertions and deletions larger than 1.5 kbp detected in the CHM1 and CHM13 homozygous cell lines, and the *in silico* CHM1/13 mixture. Sensitivity is defined as the fraction of the original CHM1 and CHM13 variants that are also detected in the mixture assembly, while positive predicted value (PPV) is defined as the proportion of the mixture’s SVs that are inherited from the original CHM1 and CHM13 assemblies. SVs detected in CHM1 only or CHM13 only are expected to be heterozygous and those detected in both are expected to be homozygous in the mixture assembly. Overall, NGM has detected a total of 2156 SVs larger than 1.5 kbp from CHM1 and CHM13, with 1370 insertions and 786 deletions. When the mixture was used as the input data, the NGM SV pipeline detected 1999 SVs: 1254 insertions and 745 deletions. This method has an overall detection sensitivity of 92.7%, with 91.5% sensitivity and 97.9% PPV for insertions, and 94.8% sensitivity and 97.1% PPV for deletions. It detected 976 homozygous and 1180 heterozygous SVs from the two haploid samples and 968 homozygous and 1031 heterozygous SVs from the pseudo-diploid mixture. In this case, it has a 99.2% sensitivity for homozygous calls and 87.4% sensitivity for heterozygous calls.

**Table 3:**
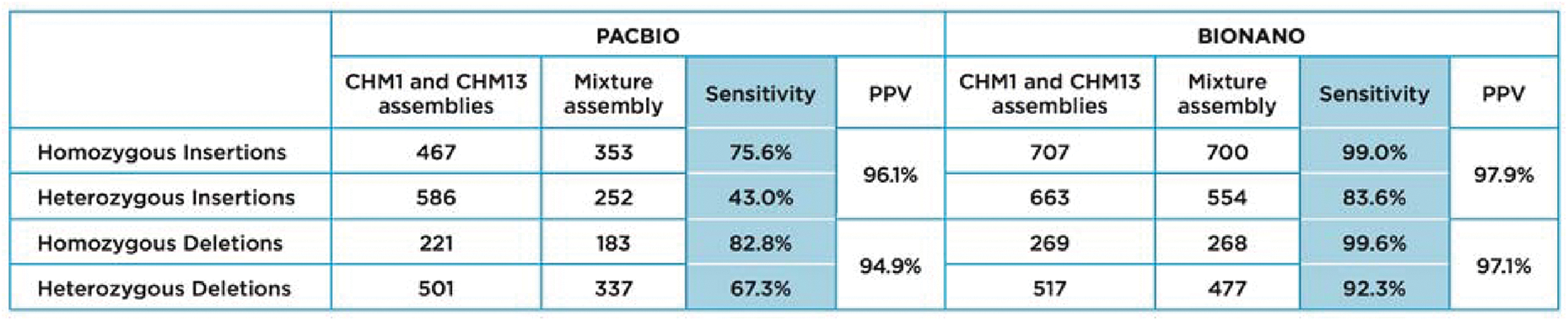
Two homozygous cell lines, CHM1 and CHM13 were independently *de novo* assembled and insertions and deletions >1.5 kbp were called. Raw data from both cell lines were combined and assembled, and SVs were called on the new assembly (mixture assemblies column). The sensitivity and PPV to detect heterozygous relative to homozygous SVs is shown.

|  | PACBIO |  |  |  | BIONANO |  |  |  |
| --- | --- | --- | --- | --- | --- | --- | --- | --- |
|  | CHM1 and CHM13 assemblies | Mixture assembly | Sensitivity | PPV | CHM1 and CHM13 assemblies | Mixture assembly | Sensitivity | PPV |
| Homozygous Insertions | 467 | 353 | 75.6% | 96.1% | 707 | 700 | 99.0% | 97.9% |
| Heterozygous Insertions | 586 | 252 | 43.0% |  | 663 | 554 | 83.6% |  |
| Homozygous Deletions | 221 | 183 | 82.8% | 94.9% | 269 | 268 | 99.6% | 97.1% |
| Heterozygous Deletions | 501 | 337 | 67.3% |  | 517 | 477 | 92.3% |  |

A similar experiment using PacBio long-read sequencing was described recently (Huddleston, 2016). Structural variants in CHM1 and CHM13 were called separately with the SMRT-SV algorithm, and compared to those called using an equal mixture of reads of both haplotypes at 60x combined coverage depth.

The reported total number of SVs larger than 1.5 kbp detected by PacBio sequencing was 1775 from CHM1 and CHM13, with 1053 insertions and 722 deletions. When the pseudo-diploid mixture was used as the input data, the reported total number of SVs was 1125, with 605 insertions and 520 deletions. This method has an overall detection sensitivity of 63.4%, with 57.5% sensitivity and 96.1% PPV for insertions, and 72.0% sensitivity and 94.9% PPV for deletions. The pipeline reported 688 homozygous and 1087 heterozygous SVs from two haploid samples, and 536 homozygous and 589 heterozygous SVs from the pseudo-diploid, resulting in sensitivities of 77.9% for homozygous and 54.2% for heterozygous SVs.

### SV performance compared to other benchmark experiments in diploid cells

Mak et al., mapped genome wide structural variation using Bionano NGM in the CEPH trio (NA12878, NA12891, NA12892) using automated and manual bioinformatics approaches. They found 7 times more large insertions and deletions (longer than 5 kbp) than previous studies based on high-depth Illumina short reads (1000 Genomes Project Consortium et al. 2010, 2012). We used our current fully automated approach to compare with that benchmark study, and found 89% of their reported SVs while also finding fourfold as many total SVs, as a result of improved sensitivity with the new assembly and SV algorithms. We have also benchmarked NGM insertion and deletion calling with PacBio SV calling as reported by Chaisson et al. for CHM1 (Chaisson MJ H. J., 2015). Based on this comparison; we show that NGM confirmed 66% of SVs called by PacBio while calling an additional 19% more SVs, comparing the total number of SVs by each technology. PacBio sequencing captures 57% of Bionano SVs. SV call sizes have a median deviation of 91 bp between Bionano and PacBio based calls (Figure 2C), which is larger than the 49 bp median error found in the *in silico* simulation, but still represents very good concordance. The increased deviation was likely caused, at least partially, by compounded error from making two empirical measurements compared to having a ground truth as in the *in silico* test.

**Figure 2:**
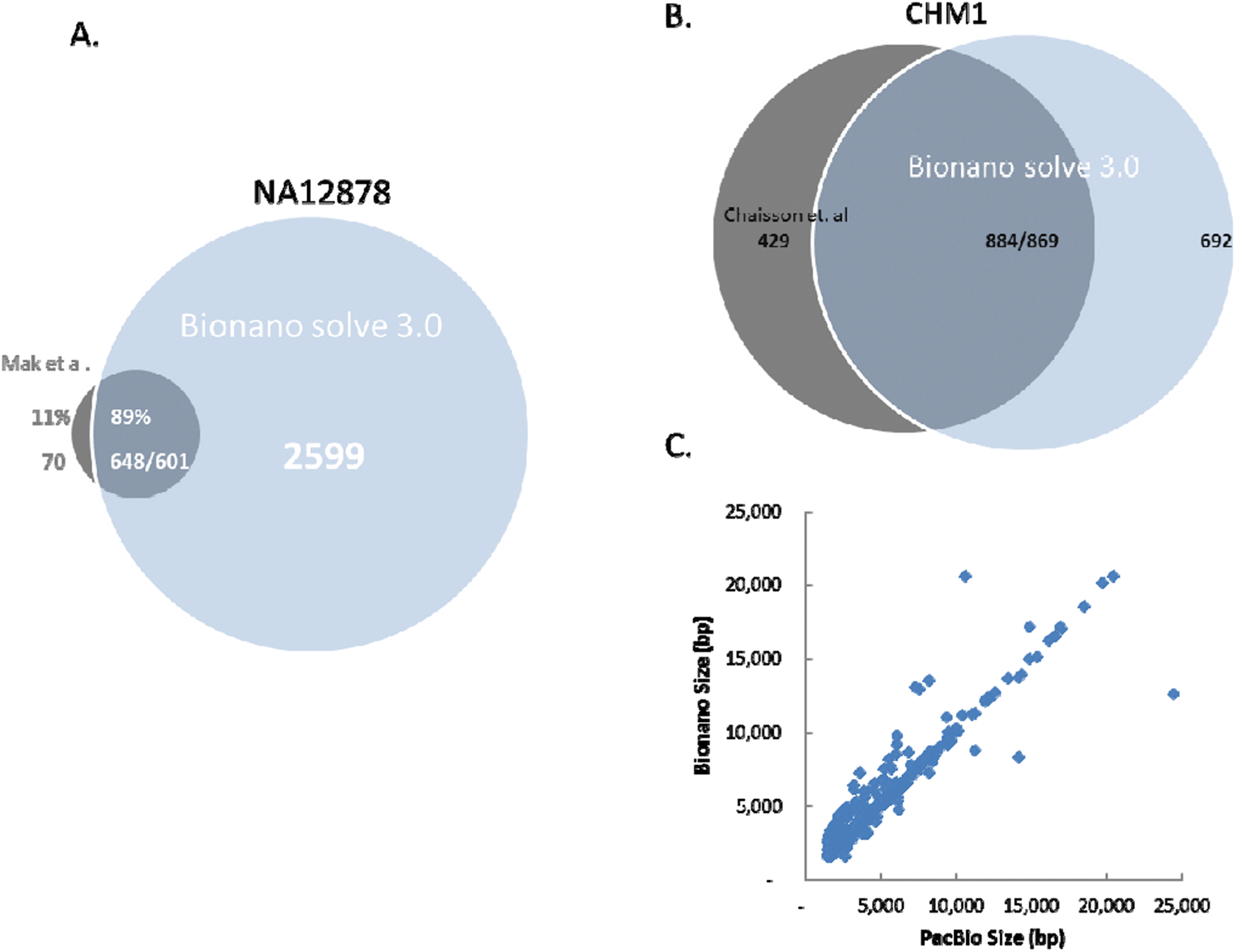
A. Comparison of insertions and deletions detected by Mak et al. through automated and manual curation, and those detected with the automated Bionano Solve 3.0 pipeline. B. Comparison of SVs detected through PacBio sequencing and SMRT-SV and those detected by Bionano Solve 3.0 for CHM1. C. Plot of the size of overlapping SVs from B.

In addition, translocation detection sensitivity was verified in two reference samples, NA16736 and NA21891, which are lymphoblast cell lines produced from blood cells from a patient with a developmental disorders resulting in deafness with DNA repair deficiency caused by t(9;22) translocation, and a second patient with Prader-Willi syndrome associated with a t(4;15) translocation, both of which have been characterized by traditional cytogenetic methods. NGM was able to detect both expected translocations as well as the reciprocal translocation breakpoints. Additionally, NA16736 contained a t(12:12) rearrangement which flanked an inverted segmental duplication. In NA21891, one translocation breakpoint could be localized within a gene, resulting in a predicted truncation (Figure 3).

**Figure 3:**
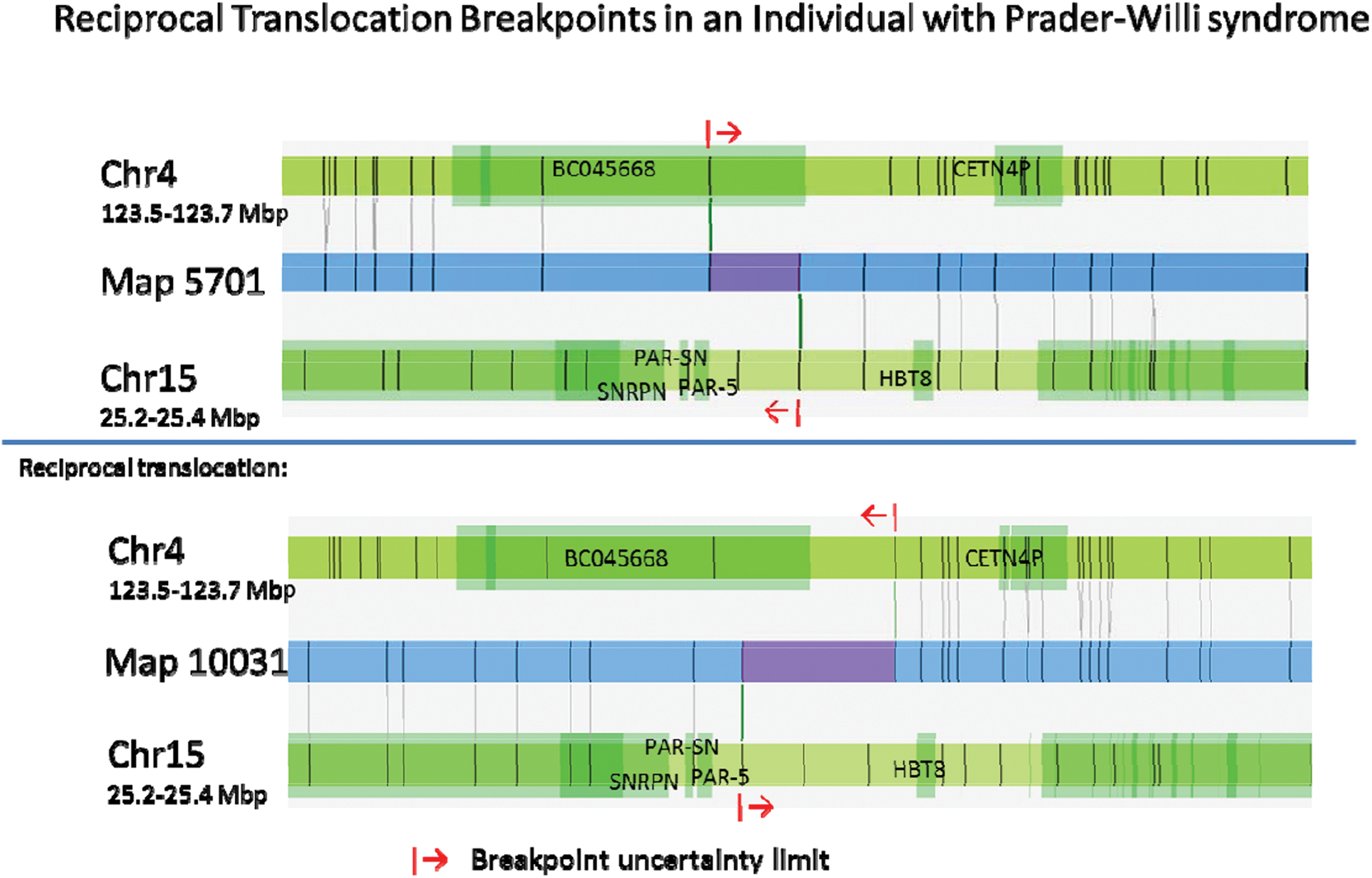
Example of a translocation detected by NGM, associated with Prader-Willi syndrome. Blue bars are Bionano maps, and vertical lines represent Nt.BspQI label sites. For each of the reciprocal translocation breakpoints, maps are shown with alignments of the maps to chromosome 4 (top) and chromosome 15 (bottom) of the human reference hg19. Breakpoint resolution can be determined by the distance between matched and unmatched labels.

## Discussion

We have tested Bionano Genomics’ NGM platform and new algorithm pipelines for sensitivity and accuracy of detection of large SVs (> 1 kbp) of *in silico* diploid genomes, as well as with several human cell lines with known structural features.

The *in silico* created diploid genome based on hg19 and an edited version of the same maps incorporating simulated SVs were used to test the SV calling capability for heterozygous variants. The artificial insertions and deletions were random sets of all sizes, mimicking empirical data, although the perfect 50:50 mixing of two artificial haploid does not always reflect the true nature of biological samples.

We found excellent overall sensitivity of over 90% for insertions and deletions > 1 kbp. We tested sensitivity for insertions and deletions from 200 bp to 1 Mbp. We have not focused on small events, as these are well addressed by NGS, especially long read sequencing.

We similarly simulated thousands of translocations and found an overall sensitivity of 98% to detect the translocation breakpoints. We correctly identified translocations and the reciprocal breakpoints in two cell lines with known translocations. Compared to traditional cytogenetics, NGM has high resolution of SV breakpoints, resulting in 2.9 kbp median accuracy for breakpoint resolution, often sufficient for PCR and sequencing if single nucleotide resolution of the fusion point is desired for subsequent gene function study.

By *de novo* assembling each of two hydatidiform moles genome (CHM1 and CHM13) separately followed by assembling a mixture of CHM1/13 molecules, we were able to compare the pseudo-diploid SV calls against the SV calls in each single haploid assembly. Huddleston et al reported a similar experiment by SV detection using SMART-SV algorithm with Pacbio sequencing data on CHM1 and CHM13 pseudodiploid genome. They have reported to identify 44% as many SVs in this single pseudodiploid genome as previously reported for the entire 2504 diploid genome in phase 3 of 1000 Genome Project and 89% of the SV called by the SMART-SV method were missed by the previous short read based genomes. The detection range for SMART-SV is from 50 bp to 28 kb, so for the purpose of a direct comparison, we only included SV calls larger than 1 kbp in this study. Here using Bionano’s SV calling algorithm the sensitivity to detect insertions and deletions > 1.5 kbp was 87.4% for heterozygous and 99.2% for homozygous variants. This far exceeds the sensitivity reported for other technologies for insertions, deletions, and translocations above 1 kbp, including the already impressive recent results by Huddleston et al. (Huddleston J, 2016).

In a detailed comparison, Bionano had 381 (21.5%) more SV calls than the PacBio based study in the haploid cell lines, and 874 (77.7%) more SV calls in the diploid genome. The sensitivity to detect homozygous SVs > 1.5 kbp in the simulated diploid experiment using PacBio was 77.9%, compared to 99.2% for Bionano. In the case of heterozygous SV calls, which is more challenging for any technology and method, the sensitivity to detect heterozygous SVs > 1.5 kbp using PacBio was 54.0%, compared to 87.4% sensitivity for Bionano. Furthermore, PacBio has only 43.0% sensitivity for heterozygous insertions and 67.3% for heterozygous deletions larger than 1.5 kbp, while notably, Bionano NGM has a much higher 83.6% detection sensitivity for heterozygous insertions and 92.3% for heterozygous deletions larger than 1.5 kbp.

We have further tested this new pipeline in a well characterized diploid cell line NA12878, which was previously mapped with Bionano Irys technology. Our new SV calling pipeline improved significantly on the published work by fully automating the detection of insertions and deletions, and by greatly improving sensitivity partially by expanding the size range of the variants down to 1 kbp from the published 5 kbp.

All current methodologies for detecting SVs have significant limitations. Chromosomal microrarray has good performance for large duplications and deletions but is insensitive to novel insertions, mobile element insertions, many low copy repeats, and all balanced translocations and inversions. Short read based methods have poor sensitivity to most large variants. Long read sequencing performs well for homozygous SVs up to around their read lengths but has relatively low sensitivity for heterozygous SVs, including smaller ones and those involving larger repetitive regions.

Precision medicine starts with a precision genome, yet obtaining complete and accurate genomic structural variation information efficiently and inexpensively remains the biggest challenge for the genomic and medical community. Bionano Genomics’ NGM is currently the most cost-effective, relatively fast, and accurate technology able to detect a wide spectrum of SV types – balanced and unbalanced, simple and complex, spanning a wide size range from 1 kbp to megabases. The sensitivity to detect heterozygous SVs reported here is superior to other methodologies for a broad range of SV types. Furthermore, NGM has a resolution that is orders of magnitude better than karyotyping and FISH. Bionano Next Generation Mapping is an essential tool for generating a complete picture of the structure of genomes, paving the way to realizing the full potential of applying genomic information towards translational research and clinical diagnostics and therapeutics.

## Materials and methods

### Isolation and fluorescent labeling of genomic DNA

For CHM1 and CHM13, genomic DNA was isolated using Bionano’s OptiDNA prep. Briefly, cells were embedded into a thin layer of low-melting point agarose, using a specialized cassette. The cells were then treated in the cassette with Puregene Proteinase K (Qiagen) and RNase A (Qiagen), resulting in purified DNA protected by an agarose matrix. For 2 hours at 37°C, DNA was digested with nicking endonucleases Nt.BspQI (New England BioLabs). Nicked DNA was then incubated for 1 hour at 50°C with Taq polymerase (New England BioLabs) and fluorescently labeled dUTP (Bionano Genomics). Next, the labeled DNA was incubated for 30 minutes at 37°C with Taq ligase (New England BioLabs) and dNTPs. The samples were then removed from the cassettes, melted, and solubilized using Agarase(Thermo Fisher Scientific), and the DNA was counterstained with YOYO-1 (Thermo Fisher Scientific).

For NA12878, NA16736, and NA21891, the standard IrysPrep was used for DNA isolation and labeling. Briefly, genomic DNA was isolated using the CHEF Mammalian Plug Kit (BioRad): after embedding cells into agarose plugs, the plugs were treated with Proteinase K Qiagen) and RNase A (Qiagen). The plugs were then melted, solubilized using Agarase (Thermo Fisher Scientific), and drop dialyzed against TE Buffer. DNA was quantitated using PicoGreen fluorescence assay. and 900 ng of DNA were labeled using the IrysPrep labeling kit. For 2 hours at 37°C, DNA was digested with nicking endonucleases Nt.BspQI (New England BioLabs). Nicked DNA was then incubated for 1 hour at 72°C with Taq polymerase (New England BioLabs) and fluorescently labeled dUTP (Bionano Genomics). Next, the labeled DNA was incubated for 30 minutes at 37°C with Taq ligase (New England BioLabs) and dNTPs. Finally, the DNA was counterstained with YOYO-1 (Thermo Fisher Scientific).

### Improvements to assembly and analysis pipeline

We introduced a number of improvements to the Bionano Solve 3.0 assembly and analysis pipeline:

Assembly: Bionano Solve 3.0 enables improved insertion, deletion, and translocation detection. While the basic strategy of overlap-layout-consensus assembly remains the same, new haplotype-aware assembly components were implemented to more effectively accommodate large heterozygous variants.

During extension stages of assembly, by analyzing molecule-to-genome map alignments, clusters of molecules with a coordinated disrupted alignment result in splitting the genome map for independent assembly. Molecules from different alleles are “peeled off”; therefore, it is possible to handle more than two alleles. This new component is critical for assembly of haplotype maps with large differences, and for detection of variants across a wide size range. It improves assembly of segmental duplication regions, where large stretches of sequence appear more than once in the genome.

*De Novo* assembly was performed using BioNano’s custom assembler software based on the Overlap-Layout-Consensus paradigm. Pairwise comparison of all DNA molecules was performed to create a layout overlap graph, which was then used to create the initial consensus genome maps. By realigning molecules to the genome maps (Refine-B P-Value 10^-11^) and by using only the best match molecules, a refinement step was performed to refine the label positions on the genome maps and to remove chimeric joins. Next, during an extension step, the software aligned molecules to genome maps (Extension P-Value 10^-11^), and extended the maps based on the molecules aligning beyond the map ends. Overlapping genome maps were then merged using a Merge P-Value cutoff of 10^-15^. These extension and merge steps were repeated five times before a final refinement was applied to “finish” all genome maps (Refine Final P-Value 10^-11^). Two assemblies were constructed per sample – one for each nicking endonuclease.

During the extension step, the software identified clusters of molecules that aligned to genome maps with unaligned ends of at least 30 kbp. If such a cluster exists, the genome map is copied, this end of it is removed and replaced by an extension-refinement of these molecules (extend-split). In addition, the final refinement step searched for clusters of molecules aligned to genome maps with internal alignment gap of size < 50 kbp, in which case, the genome maps were changed into two haplotype maps. The extend-and-split function is essential to identify large allelic differences and to assemble across loci with segmental duplications, whereas the refinement haplotype function can find smaller differences.

#### SV identification

SVs are detected based on analyzing two or more local alignments between the de novo assembled genomes and a public human reference assembly. We required an alignment cutoff of P-Value of 10^-12^. SV calling was done for the Nt.BspQI and Nb.BssSI assemblies independently. Significant discrepancies in the distance between adjacent labels or the number of unaligned labels between adjacent aligned labels (outlier P-Value 3x10^-3^) indicated the presence of insertion and deletions. Translocation breakpoints are detected as fusion points between supposedly distant regions of the genome. Intrachromosomal translocation breakpoints involve regions at least 5 Mbp apart. Interchromosomal translocation breakpoints by definition involve regions on different chromosomes.

#### SVMerge

SVMerge for insertions, deletions, and translocations allows users to take advantage of having two single-enzyme datasets (for example, Nt.BspQI and Nb.BssSI). SVMerge provides several potential benefits. The complementary nature of the two enzymes helps improve sensitivity compared to one-enzyme detection. Cross-confirmation by two independent assemblies provides a useful means of validating variant calls. We expect improved SV breakpoint accuracy due to higher combined labeling site density. For insertions and deletions, SV size estimates are expected to be more accurate for the same reason. Confidence scores for insertions and deletions reflect whether just one or both enzymes support an SV call.

Briefly, SVMerge examines SV calls from single-enzyme assemblies and evaluates whether they are overlapping calls. Overlapping calls are then merged, and refined breakpoint coordinates are output.

### Construction of simulated diploid genomes

We simulated random SV events so that we could estimate our genome-wide SV calling performance accurately. The human reference assembly hg19 was used as an “SV-free” base genome. Random SV events were simulated to form an edited genome.

An initial total of 1600 insertions and 1600 deletions were randomly introduced into an *in silico* map of the human reference genome, hg19, such that indels ranged from 200 bp to 1 Mbp, Indels were separated by at least 500 kbp.

Two types of translocation events were simulated. In “cut-and-paste” genomes, an initial of 1,000 segments were randomly selected across hg19 and randomly inserted elsewhere in the genome, creating 3,000 breakpoints and 2,000 translocation events to be detected by the pipeline. N-base gaps were avoided. The size of the translocation fragments ranged from 50 kbp to 1 Mbp. Translocation breakpoints were at least 500 kbp away from each other. For intrachromosomal translocations, the breakpoints were at least 5 Mbp apart. Two types of reciprocal translocations were simulated. In genome 1, two breakpoints were randomly simulated on each chromosome, and the ends of the chromosomes were reciprocally exchanged between them. In genome 2 and 3, translocation events from 6 published studies were simulated. In all types of translocations, the genomic material remained the same but with a different arrangement compared to the unmodified hg19. The exchange is balanced, and thus no copy number variation was generated.

Once the edited genomes were constructed, molecules were simulated by carving randomly along the genome. Two sets of molecules were simulated, each based on a different nicking endonuclease. Based on the edited and the unedited hg19, molecules were simulated to resemble actual molecules collected by Bionano NGM technology. Errors were added to molecules to mimic the uncertainty of DNA and endonuclease internal properties, sample preparation, and data collection equipment. The same was done for the hg19 genome. Then molecules from these two genomes were mixed to form a diploid genome, where all SV events are heterozygotes.

Simulated molecule datasets, with 70x effective coverage, were generated and assembled and used as input for the Bionano Solve 3.0 pipeline. SV calls were made by combining the data from both nick motifs such that matching SV calls from two enzymes were merged into a single call. The final merged SV calls were compared to the *in silico* ground truth SV call set.

## Acknowledgments

We thank Saki Chan for technical and proofreading assistance, Khoa Pham and Jeff Reifenberger for technical help, Pui Yan Kwok and Evan Eichler for helpful discussions, and Carl Baker for providing cell cultures for CHM1 and CHM13.

